# The ecological significance of extracellular vesicles in modulating host-virus interactions during algal blooms

**DOI:** 10.1101/2021.02.10.430559

**Authors:** Daniella Schatz, Guy Schleyer, Marius R. Saltvedt, Ruth-Anne Sandaa, Ester Feldmesser, Assaf Vardi

**Author notes:** Corresponding author: Assaf Vardi.

## Abstract

Extracellular vesicles are produced by organisms from all kingdoms and serve a myriad of functions, many of which involve cell-cell signaling, especially during stress conditions and host-pathogen interactions. In the marine environment, communication between microorganisms can shape trophic level interactions and population succession, yet we know very little about the involvement of vesicles in these processes. In a previous study, we showed that vesicles produced during viral infection by the ecologically important model alga *Emiliania huxleyi*, could act as a pro-viral signal, by expediting infection and enhancing the half-life of the virus in the extracellular milieu. Here, we expand our laboratory findings and show the effect of vesicles on natural populations of *E. huxleyi* in a mesocosm setting. We profile the small-RNA (sRNA) cargo of vesicles that were produced by *E. huxleyi* during bloom succession, and show that vesicles applied to natural assemblages expedite viral infection and prolong the half-life of this major mortality agent of *E. huxleyi*. We subsequently reveal that exposure of the natural assemblage to *E. huxleyi*-derived vesicles modulates not only host-virus dynamics, but also other components of the microbial food webs, thus emphasizing the importance of extracellular vesicles to microbial interactions in the marine environment.

## Main Text

The eukaryotic phytoplankter *Emiliania huxleyi* is a major contributor to the marine ecosystem. In temperate oceans, *E. huxleyi* forms massive blooms that are the basis of the marine food web and greatly influence the biogeochemical cycles of carbon and sulfur (1, 2). Annual blooms of *E. huxleyi* are frequently infected by a large dsDNA virus, the *E. huxleyi* virus (EhV) (3). Like all other organisms, *E. huxleyi* produces extracellular vesicles (hereafter, vesicles) that can serve as a mode of communication between the producing cells and specific target cells (4, 5). We previously showed that cultured *E. huxleyi* increases the production of vesicles during viral infection. The lipid composition of these vesicles is signifcantly different to that of their producing cells, and is composed mainly of triacylglycerols. As cargo, the vesicles carry small RNAs (sRNAs) that are predicted to target host cell cycle and lipid metabolism genes. Vesicles derived from EhV-infected cultures expedited viral infection and prolong the half-life of EhV in a dose dependent manner (6).

We sought to establish the ecological role of vesicles in the natural environment, and specifically, their effect on algal bloom demise and subsequent succession. To this aim, we set up a mesocosm experiment at the Marine Biological Station in Espegrend, Norway. The 23-day experiment included four mesocosm bags that contained the natural microbial assemblage. A bloom of *E. huxleyi* was induced (Figure 1a) by adding nitrogen and phosphorus to the water ((7), see methods). During the course of bloom sucession, we identified three major phases: pre-bloom (days 1-9), bloom (days 10-17) and demise (days 18-23). In the demise phase, we observed an increase in the abundance of EhV in the water based on qPCR quantification of viral *mcp* (Figure 1b), which suggests that the termination of the bloom, at least in part, was due to viral infection (8).

**Figure 1.**
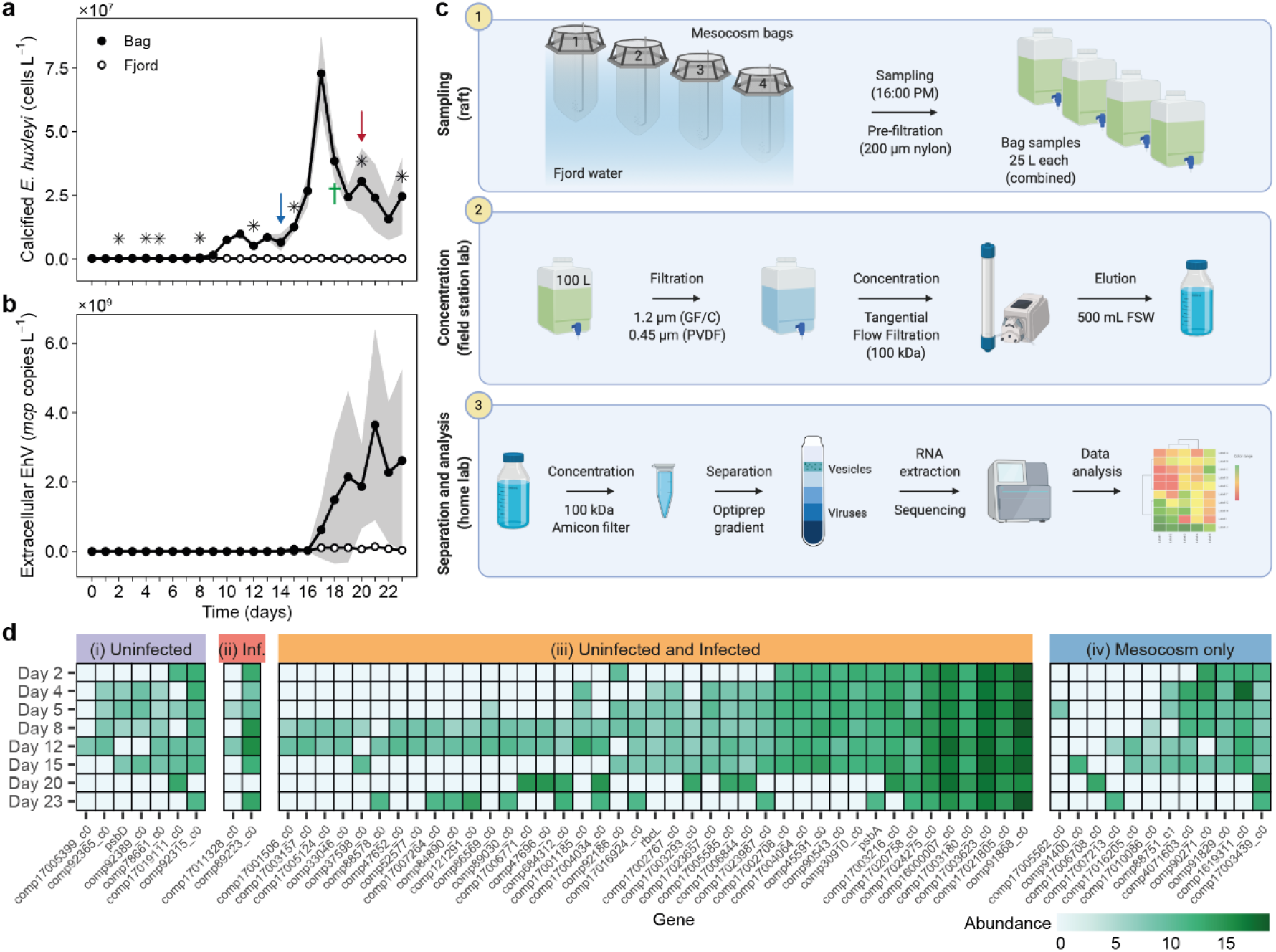
Extracellular vesicles from natural *E. huxleyi* populations. A bloom of *E. huxleyi* was induced in a mesocosm setup at the Marine Biological Station Espegrend, Norway (60°16′11N; 5°13′07E) in May-June 2018. Abundance of calcified *E. huxleyi* cells (a) was measured by flow cytometry, and EhV concentration (b) was measured by qPCR targeting the major capsid protein gene (*mcp*). Average +/− SE of the four mesocosm bags are presented for (a) and of bags 2 and 4 in (b). Abundance of cells and virions was also measured in the surrounding fjord water as a “blank” control (empty circles). Asterisks indicate the times at which samples were taken for vesicle extraction; arrows indicate sampling times for the experiments presented in Figure 2a, b; cross indicates time of sampling of EhV for the experiment presented in Figure 2c. (c) Workflow for vesicle sRNA profiling. Sampling (1): 25 l water samples were collected on the days marked with an asterisk in (a) from each bag using a peristaltic pump and a 200 µm nylon pre-filter. Concentration (2): the water samples were combined and filtered through two subsequent filters (GF/C and 0.45 µm PVDF). Samples were then concentrated to ~500 ml on a 100 kDa TFF cartidge and stored at 4^°^C in the dark. Separation and analysis (3): Once at the home lab, the samples were further concentrated on 100 kDa Amicon-ultra filters, and separated on an Optiprep gradient. After speparation and cleaning, vesicles were subjected to RNase treatment to eliminate extra-vesicular RNA. sRNA within the vesicles was then extracted and sequenced. Workflow was created with BioRender.com. (d) Vesicles were collected from the natural assemblages at the time points indicated by an asterisk in (a) and the sRNA cargo was sequenced. sRNA sequences were aligned to *E. huxleyi* target genes as indicated. sRNAs that target the same genes were also found in vesicles from lab cultures of (i) uninfected *E. huxleyi* CCMP2090, (ii) infected cultures, (iii) Both uninfected and infected cultures or (iv) not found in vesicles from lab cultures of *E. huxleyi* CCMP2090. Read counts were scaled to one million reads mapped to the *E. huxleyi* transcriptome, log2 transformed and compared across time points.

Throughout the bloom we collected and isolated vesicles from the mesocosm bags by concentrating them with tangential flow filtration (TFF) and enriching them by gradient separation (Figure 1c). In a previous study we showed that the main cargo of *E. huxleyi*-derived vesicles is small RNA (6). Therefore, we constructed and sequenced sRNA libraries from the vesicles isolated during the bloom. By mapping these sequences to the *E. huxleyi* transcriptome (9), we confirmed that some of these vesicles were produced by *E. huxleyi*. This was further supported by comparing the sequences to the sRNA sequences found in vesicles from lab-based cultures. We compared the potential target genes of the sRNA sequences from the natural vesicles to those of vesicles from laboratory cultures of *E. huxleyi*, and devided them into four categories based on the their presence in vesicles from lab cultures (Figure 1d): i) Seven potential target genes that were identified in cargo of vesicles from uninfected lab cultures; ii) Two potential target genes that were identified in vesicles from lab cultures infected with EhV; iii) 41 target genes that were identified to in the cargo of vesicles from both infected and uninfected cultures and iv) sequences of twelve genes that were identified in the cargo of natural vesicles, but were not found in vesicles from lab cultures. Most of the sRNA-target genes that we detected are found in multiple samples, which may suggest that they are ubiquitus to vesicles from *E. huxleyi* blooms. We also identified sRNA sequences that potentially target genes that were not targeted by sRNA in lab-based vesicles. This could be due to secretion of vesicles from *E. huxleyi* cells exposed to varying environmental conditions or during interactions of *E. huxleyi* with other microorganisms (e.g. bacteria) (10), that induces packaging of different sRNA molecules into vesicles. Aditionally, strain-specific sRNA cargo packaging can explain why these sequences were not identified in lab-based vesicles. Interestingly, despite the strong evidence for viral infection (Fig. 1b), we could only find two sequences that are related to viral infection. One of these sequences was abundant throughout the experiment, even in the pre-bloom stage.

The sRNA in vesicles may regulate specific genes in target *E. huxleyi* cells (Figure 1d), and may coordinate, to some extent, their metabolic state. For example, in vesicle samples from all time points, we detected sRNA sequences that potentially regulate the *E. huxleyi* gene comp17003623_c0, which encodes a putative cation transport protein (Table S1). Thus, these cargo molecules may change the ability of target *E. huxleyi* cells to transport organic molecules and, consequently, change their metabolic profile. In addition, we found sRNA sequences that potentially regulate *hsp*70 that is involved in stress response, including high light intensity (11). We also detected the presence of sRNA of *rbc*L and *psb*A that encode the RubisCO large subunit and the photosystem II D1 protein, respectively, suggesting a potential regulation of photosynthesis by vesicles.

In order to examine the biological effect of vesicles on infection dynamics during algal blooms, we exposed natural assemblages from the bloom to vesicles isolated from lab cultures of *E. huxleyi* CCMP2090 at a ratio of ~500 vesicles per *E. huxleyi* cell, which is the estimated ratio in lab cultures at the end of infection (6). When applied to microbial communities from the early bloom phase (day 14), the vesicles had little to no effect on the growth of major phytoplankton groups (Figure 2a). We observed a small reduction in bacterial abundance 48h after vesicle treatment, which suggests that the bacteria do not benefit from the vesicles and do not use them as a source for nutrients, as suggested for vesicles of *Prochlorococcus* (12). Interestingly, treatment of natural population taken from *E. huxleyi* demise phase (day 20, Figure 2b), when EhV was clearly present in the population (Figure 1b), led to a significant reduction in the abundance of nanophytoplankton, a group that includes *E. huxleyi*. This was accompanied by an elevated EhV abundance as compared to the control populations that were not exposed to vesicles (Figure 2b). We suggest that extracellular vesicles expedite viral infection not only in *E. huxleyi* cultures but also during bloom demise. We did not observe an effect of vesicles on the calcified *E. huxleyi* population which could be due to simultaneous growth of virus-resistant strains (13) or induction of a non-lytic infection stage (8, 14). This experimental setup enabled us to examine the effect of vesicles from *E. huxleyi* on other members of the microbial food web. Indeed, we observed an elevated abundance of *Synechococcus* in the vesicle-treated samples, compared to those that were not exposed to vesicles. This result accentuates the effect of *E. huxleyi*-produced vesicles, not only on *E. huxleyi* itself, but also on other photosynthetic microorganisms in the natural assemblage. We propose that *E. huxleyi* and *Synechococcus* compete for resources, and when vesicles expedite the viral infection of *E. huxleyi, Synechococcus* can grow faster. An alternative scenario can be that allelopathic interactions between *E. huxleyi* and *Synechococcus* may suppress *Synechococcus* growth during the *E. huxleyi* bloom. When vesicles are plentiful, viral infection of *E. huxleyi* is expedited and the allelopathic pressure is potentially alleviated from *Synechococcus*. This result enables the expansion of the impact of the viral shunt during algal bloom demise from heterotrophic (15) to photosynthetic bacteria.

**Figure 2.**
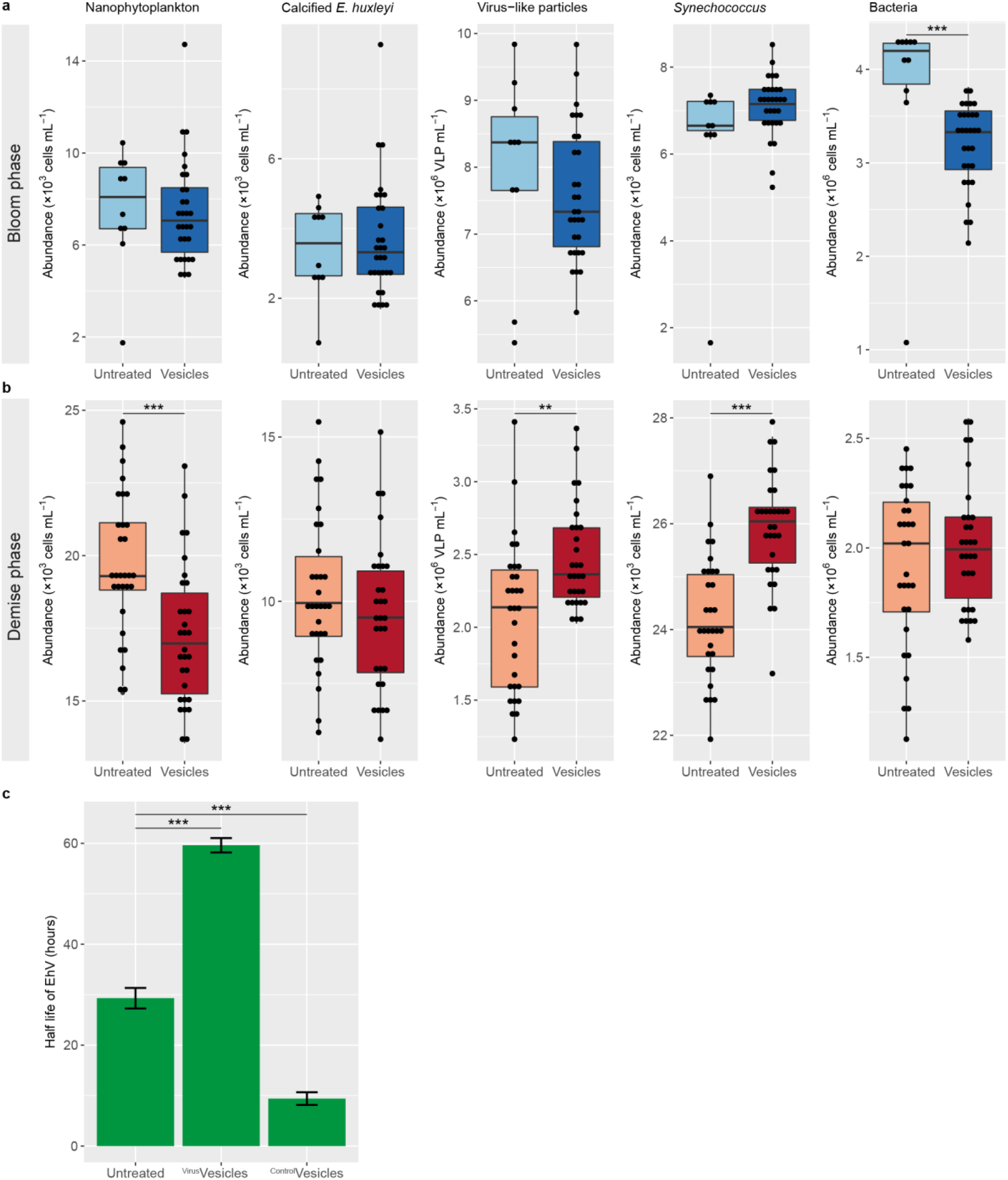
Extracellular Vesicles modulate viral infection of *E. huxleyi* and prolong the half-life of EhV in natural assemblages. Vesicles derived from lab cultures of *E. huxleyi* CCMP2090 were incubated with natural microbial populations from (a) the early bloom phase (day 14, blue arrow in Figure 1a) or (b) demise phase (day 20, red arrow in Figure 1a). Abundance of nanophytoplankton (including calcified and non-calcified *E. huxleyi*), calcified *E. huxleyi*, EhV-like particles, *Synechococcus* and bacteria was measured by flow cytometry 48h post-treatment. In (a), significant difference between the treated and untreated populations was observed only for bacteria (\*\*\**p* value = 0.001). For untreated *n* = 10, for vesicle-treated *n* = 30. In (b), significant reduction in nanophytoplankton abundance (\*\*\**p* value = 0.0003), concomitant with elevation in EhV-like particle count (\*\**p* value = 0.009) and *Synechococcus* abundance (\*\*\**p* value = 1.03 × 10^-7^) was observed. For both treated and untreated *n* = 30. At least 10,000 events were counted for each gate. In (a) and (b), *p* value was calculated using two tailed t-test with equal variance. (c) Half-life of infectious EhV from the mesocosm bags, measured by the most probable number (MPN) method. EhV was sampled from the bags during the demise phase of the bloom (day 18, green cross in Figure 1a). Vesicles from infected (^Virus^Vesicles) or uninfected (^Control^Vesicles) lab cultures of *E. huxleyi* CCMP2090 were added to natural EhV at a ratio of 10 vesicles per EhV-like particle. For the ^Control^vesicles treatment, the decay was so fast we could not detect infectivity in more than one time point. Therefore, the minimum detectable infectivity values were used in the subsequent time points in order to calculate the maximum possible half-life. Average +/− SE is presented. \*\*\**p* value<0.001 for each treatment compared to the untreated control, using ANOVA with Dunnett’s post-hoc test, *n* = 3.

We previously showed that under laboratory conditions, incubating EhV virions with vesicles derived from infected cultures led to a prolonged half-life of the viruses (6). To test the ecological significance of this phenomenon, we incubated natural EhV with vesicles derived from both infected and uninfected lab cultures of *E. huxleyi* under natural light and temperature conditions, and examined the decay by assessing the loss of infectivity over time (Figure 2c). Vesicles derived from infected *E. huxleyi* cultures led to a prolonged half-life of natural EhV, in which we observed a 50% decay in ~60 h, as compared to ~30 h in the untreated samples. Since EhV is a specialized virus that can only infect *E. huxleyi*, this effect can have a major consequence to the survival of infective virions in the environment, where they are exposed to UV and other damaging conditions (16, 17). Interestingly, vesicles from uninfected lab cultures of *E. huxleyi* led to rapid decay of EhV infectivity. This profound effect resulted in a complete loss of infectivity after 48 h of incubation with EhV virions (Figure 2c, see Methods section).

Taken together, we propose that in the natural environment, uninfected *E. huxleyi* cells protect themselves by secreting vesicles that reduce the half-life of EhV particles (Figure 2c). However, once infection of the *E. huxleyi* population is initiated, a cascade of cellular events ultimately leads to expedited viral infection (Figure 2b), concomitant with elevation of the half-life of the virions in the environment (Figure 2c). This might have an effect on other phytoplankton species, such as *Synechococcus*, thus affecting the succession of species in the marine environment. We propose that communication via extracellular vesicles during microbial interactions in algal blooms may have a profound effect on the growth and composition of the associated marine microbial food webs.

## Materials and methods

### Mesocosm setup

The mesocosm experiment AQUACOSM VIMS-Ehux was carried out between 24^th^ May (day 0) and 16^th^ June (day 23) 2018 in Raunefjorden at the Marine Biological Station Espegrend, Norway (60°16′11N; 5°13′07E) as previously described (7). Four light-transparent enclosure bags were filled with surrounding fjord water (day −1; pumped from 5 m depth) and continuously mixed by aeration (from day 0 onwards). Each bag was supplemented with nutrients at a nitrogen to phosphorous ratio of 16:1 (1.6 µM NaNO3 and 0.1 µM KH2PO4 final concentration) on days 0-5 and 14-17, whereas on days 6, 7 and 13 only nitrogen was added. Nutrient concentrations were measured daily (18).

### Enumeration of phytoplankton cells by flow cytometry

For *E. huxleyi* enumeration by flow cytometry, water samples were collected in 50 mL tubes from approximately 1 m depth. Water samples were pre-filtered using 40 µm cell strainers and immediately analyzed with an Eclipse iCyt flow cytometer (Sony Biotechology, Champaign, IL, USA) as previously described (19). A total volume of 300 µl with a flow rate of 150 µl min^-1^ was analyzed. A threshold was applied on the forward scatter to reduce background noise. Four groups of phytoplankton populations were identified in distinct gates by plotting the autofluorescence of chlorophyll (em: 663-737 nm) versus phycoerythrin (em: 570-620 nm) and side scatter: calcified *E. huxleyi* (high chlorophyll and high side scatter), *Synechococcus* (high phycoerythrin), nanophytoplankton including calcified and non-calcified *E. huxleyi* (high chlorophyll and phycoerythrin) and picophytoplankton (low chlorophyll and low phycoerythrin) (20). See Figure S1 for further details of gating strategy.

### Enumeration of EhV-like particles and bacteria by flow cytometry

For EhV and bacteria counts, 200 µl of sample were fixed a final concentration of 0.5% glutaraldehyde for one hour at 4°C and flash frozen in liquid nitrogen. For analysis, they were thawed and stained with SYBR gold (Invitrogen, Carlsbad, CA, USA) that was diluted 1:10000 in 0.2μm filtered TE buffer (10:1 mM Tris:EDTA, pH 8), incubated for 20 min at 80°C and cooled to room temperature (21). Bacteria and EhV-like particles were counted and analyzed using an Eclipse iCyt flow cytometer (ex: 488 nm, em: 500–550 nm) and identified comparing to reference samples containing fixed EhV201 and bacteria from lab cultures.

### Enumeration of extracellular EhV by qPCR

Water samples (1-2 l) were sequentially filtered by vacuum through polycarbonate filters with a pore size of 20 µm (47 mm; Sterlitech, Kent, WA, US), then 2 µm (Isopore, 47 mm; Merck Millipore, Cork, Ireland), and finally 0.22 µm (Isopore, 47 mm; Merck Millipore). Filters were immediately flash-frozen in liquid nitrogen and stored at −80°C until further processing. DNA was extracted from the 0.22 µm filters using the DNeasy PowerWater kit (Qiagen, Hilden, Germany) according to the manufacturer’s instructions. Each sample was diluted 100 times, and 1 µl was then used for qPCR analysis. EhV abundance was determined by qPCR for the major capsid protein (*mcp*) gene (22) using the following primers: 5′-acgcaccctcaatgtatggaagg-3′ (mcp1F, (23)) and 5′-rtscrgccaactcagcagtcgt-3′ (mcp94Rv; Mayers, K. *et al*., unpublished). All reactions were carried out in technical triplicates. For all reactions, Platinum SYBER Green qPCR SuperMix-UDG with ROX (Invitrogen) was used as described by the manufacturer. Reactions were performed on a QuantStudio 5 Real-Time PCR System equipped with the QuantStudio Design and Analysis Software version 1.5.1 (Applied Biosystems, Foster City, CA, USA) as follows: 50°C for 2 min, 95°C for 5 min, 40 cycles of 95°C for 15 s, and 60°C for 30 s. Results were calibrated against serial dilutions of EhV201 DNA at known concentrations, enabling exact enumeration of viruses. Samples showing multiple peaks in melting curve analysis or peaks that were not corresponding to the standard curves were omitted.

### Vesicle concentration and separation

#### Lab samples

*E. huxleyi* CCMP2090 was grown in 20 l filtered sea water (FSW) supplemented with K/2 nutrient mix at 18°C, 16:8 h light:dark cycle, 100 μmol photons m^−2^ s^−1^. Uninfected cultures were grown to ~10^6^ cells ml^-1^. Infected cultures were inoculated with EhV201 at a multiplicity of infection (MOI) of ~1:1 plaque forming unit (pfu) per cell and incubated under normal growth conditions for 120 h, at which time the culture had cleared. The entire 20 l volume was then filtered through a GF/C filter (Whatman, Maidstone, United Kingdom) and 0.45 µm PVDF filter (Durapore, Merck Millipore) to eliminate cells and cellular debris.

#### Mesocosm samples

On days 2, 4, 5, 8, 12, 15, 18 and 23 we collected 25 l from bags 1-4 and combined them to produce one sample of 100 l for each sampling time. The samples were pre-filtered using a 200 µm nylon mesh, and then filtered through a GF/C filter (Whatman) and 0.45 µm PVDF filter (Durapore, Merck Millipore) to eliminate cells and cellular debris.

#### Particle concentration

Particles in the flow-through from the filtration stage were concentrated on a 100 kDa tangential flow filter (Spectrumlabs, Repligen, Waltham, Massachusetts, USA) to a final volume of ~500 ml. At this stage, mesocosm samples were stored in the dark at +4°C and shipped back to the home lab. All samples were further concentrated to a final volume of 1-2 ml using 100 kDa Amicon-ultra filters (Merck Millipore).

#### Vesicle separation

Vesicles were separated from other particles (including viruses) using an 18– 35% OptiPrep gradient (MilliporeSigma, St. Louis, Missouri, USA). Gradients were centrifuged in an ultracentrifuge for 12 h at 200,000×g. Fractions (0.5 ml) were collected from the top of the gradient and the fraction material was cleaned by washing three times and resuspended in 0.02 µm-filtered FSW using 100 kDa Amicon-ultra filters (Merck Millipore). Vesicles were detected in fractions with densities of 1.05–1.07 g ml^−1^ (fractions 3-5 from the top).

Vesicle concentration in samples from lab cultures was measured by NTA using the NanoSight NS300 instrument (Malvern Instruments, Malvern, UK) equipped with a 488 nm laser module and NTA V3.2 software. Samples were diluted so that each field of view contained 20-100 particles. Three 60 s videos were recorded for each biological replicate, representing different fields of view. All the videos for a given experiment were processed using identical settings (screen gain of one and detection threshold of five).

### RNA extraction and sequencing

In order to eliminate RNA molecules that are not packed into vesicles, we subjected vesicle samples to RNase treatment prior to RNA extraction. Samples were incubated for 60 min at 37°C with 10 pg µl^-1^ of RNase A (Bio Basic, Toronto, Canada). RNase activity was inactivated by adding 100 unites of Protector RNase Inhibitor (Roche, Basel, Switzerland). Total RNA (including RNA from approximately 18 nucleotides or more) was extracted using the miRNeasy kit according to the manufacturer’s instructions (Qiagen). Libraries were prepared using the TruSeq Small RNA Library kit (Illumina), according to the manufacturer’s protocol. Each sample was indexed twice with the same index, one with PNK treatment (according to manufacturer’s instructions, NEB, Ipswich, Massachusetts, USA) and one without. After 15 cycles of PCR amplification, libraries were cleaned with the QIAquick PCR Purification Kit according to the manufacturer’s instructions (Qiagen). Libraries were sequenced on the Illumina NextSeq platform.

### sRNA bioinformatics analysis

Low-quality read ends were trimmed and adaptors were removed using the cutadapt program (24), version 1.18. Reads shorter than 17 bp after the trimming were removed from further analyses. The remaining reads were mapped to an *E. huxleyi* integrated reference transcriptome shortly described in (6) using the RSEM software (25), version 1.3.1, with the default option of bowtie, version 1.1.2 (26). Genes that had at least 5 reads in any of the samples were selected. For the heatmap (Figure 1d), read counts were scaled to one million reads mapped to the *E. huxleyi* transcriptome and log2 transformed.

### Data availability

All the small RNA sequences were submitted to the SRA under BioProject PRJNA694552, accession SRR13648434 - SRR13648443.

### Effect of vesicles on natural populations – experimental design and analysis

On days 14 and 20 of the mesocosm experiment (blue and red arrows in Figure 1a, respectively), we combined equal volumes of water samples from bags 1-4 and filtered them through a 10 µm nylon mesh to eliminate zooplankton predators. We then supplemented the natural populations with f/50 nutrient mix and divided them into flasks, each containing 10 ml. 30 flasks were treated with vesicles from uninfected lab cultures of *E. huxleyi* CCMP2090, at a ratio of ~500 vesicles cell^-1^ (calcified *E. huxleyi* determined by flow cytometry), and then all flasks were incubated in a growth chamber (15°C, 16:8 h light:dark cycle, 100 μmol photons m^−2^ s^−1^). Once a day, samples were taken for flow cytometry for quantification of live cells (see “Enumeration of phytoplankton cells by flow cytometry” above), or fixed for virus and bacteria counts (see “Enumeration of EhV-like particles and bacteria by flow cytometry” above). For statistical analysis, we used two-tailed t-test with equal variance.

### Decay rate of EhV virions-experimental design and analysis

To determine the decay rate of infectivity of natural EhV virions, water were sampled from bag 4 on day 18, at a time point when viral infection was detected (green cross in Figure 1a). This sample was filtered through an 0.45 PVDF filter (Durapore, Merck Millipore) to eliminate algal and most bacteria cells. EhV-like particles were counted by flow cytometry as described above and divided into nine tubes, each containing 1 ml. Triplicate samples were either treated with vesicles from EhV201-infected (^Virus^Vesicles) or uninfected (^control^Vesicles) lab cultures (see above) at a ratio of ten vesicles per EhV-like particle, or not treated at all. All tubes were incubated in an on-land mesocosm facility that mimics the light and temperature conditions fount at ~1 m depth within the fjord water. We used the most probable number (MPN) method (27) to determine the half-life of EhV within these samples. Briefly, a series of five-fold dilutions was prepared for each sample. Each dilution (10 μl) was then added, in eight technical replicates, to 100 μl of exponentially growing *E. huxleyi* CCMP374 cultures in multi-well plates and incubated under normal growth conditions for five days. This was repeated for four consecutive days for all samples. Clearance (infection) of the cells in the multi-wells was measured using an EnSpireTM 2300 Multilabel Reader (PerkinElmer, Turku, Finland) set to *in vivo* fluorescence (ex:460nm, em:680nm). MPN was calculated using the MPN calculation program, version 5 (28). For the samples treated with ^control^vesicles, we could only obtain a positive MPN value for one time point, as the decay was faster than expected. Therefore, the minimum detectable infectivity values were used in order to calculate the maximum possible half-life. For statistical analysis, each treatment was compared to the untreated control, using ANOVA with Dunnett’s post-hoc test.

## Acknowledgments

We thank all team members of the VIMS-Ehux project for setting up and conducting the mescosom experiment, especially Flora Vincent who performed the live flow cytometry counts on the mesocosm populations, Jorun Egge, Aud Larsen and Tatiana Tsagaraki. We are grateful to Ron Rotkopf for his advice on statistical analysis. This research was supported by the European Research Council CoG (VIROCELLSPHERE grant no. 681715) and a research grants from the Estate of Bernard Berkowitz and the *de Botton* Center for Marine Science awarded to A.V. The mesocosm experiment VIMS-Ehux was supported by EU Horizon2020-INFRAIA project AQUACOSM (grant no. 731065).

## Author contributions

D.S. and A.V. conceptualized the project and wrote the manuscript. D.S and G.S. designed and performed the experiments. M.R.S. and R-A.S. performed the MPN analyses. E.F. performed all bioinformatic analyses on the RNA sequences. D.S. and G.S. extracted DNA and performed the qPCR analysis. All authors commented on the manuscript.

---

**Figure S1.**
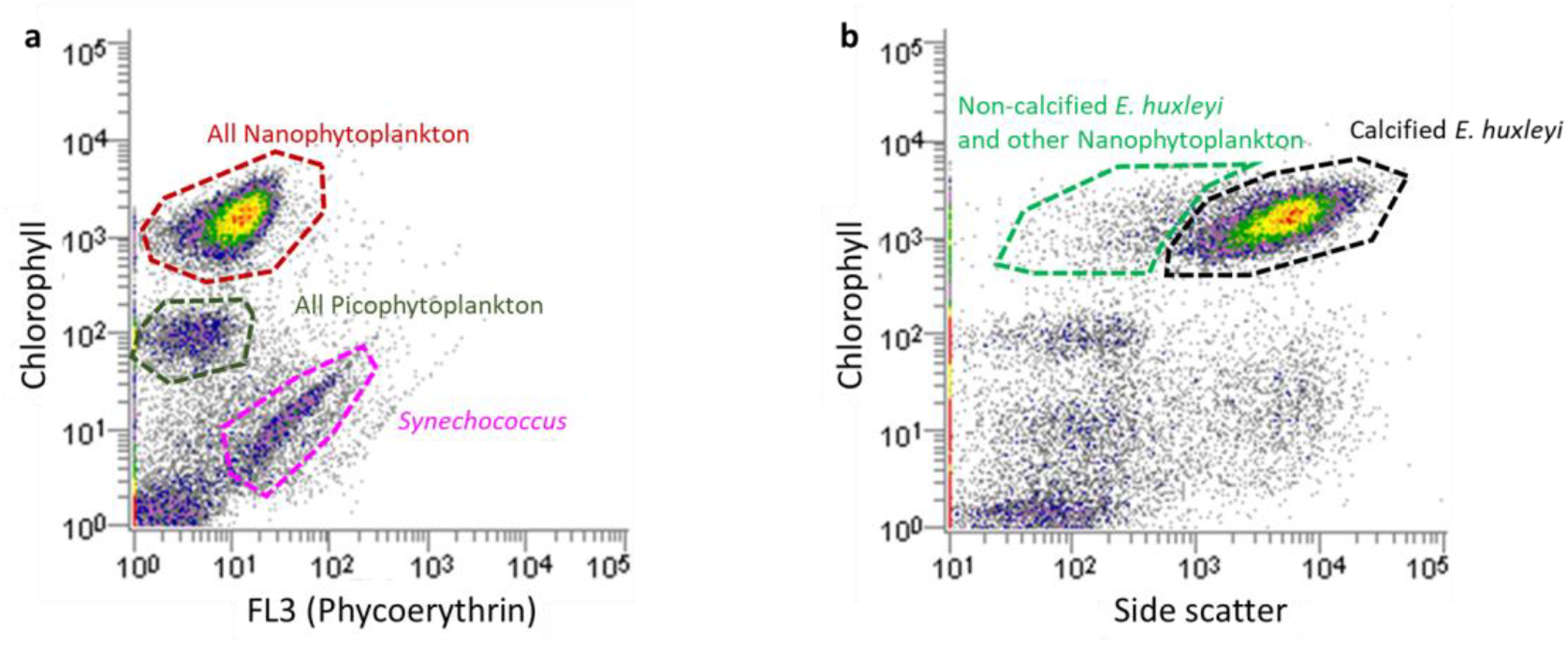
Gating strategy for flow cytometry analysis of natural phytoplankton communities. (a) Example cytogram that contains a mixture of Nanophytoplankton (red), Picoephytoplankton (green) and Synechococcus (pink). Populations were identified based on their different chlorophyll and phycoerythrin content. (b) Calcified E. huxleyi was identified based on the high side scatter caused by the coccoliths that adorn the cells.

